# Genomic Epidemiology of Canine Distemper Virus (CDV) in Neotropical Primates in Brazil Reveals Multiple Spillover Events and Convergent Evolution at Hemagglutinin Residue 519

**DOI:** 10.64898/2026.09.01.748675

**Authors:** Mayara Bertanhe, Milena G. Cabral, Ian N. Valença, Camila L. Nacif, Ingra M. Claro, Lucas A. Diniz, Maiara M. M. Valeriano, Lívia A. Gomes, Diogo H. M. Costa, Ramon S. Oliveira, Maria E. Gonçalves-dos-Santos, Esmenia C. Rocha, Bianca C. Silva, Franciane M. de Oliveira, Lucas A. M. Franco, Filipe V. S. de Abreu, Thallyta M. Vieira, Ester C. Sabino, Nuno R. Faria, Filipe R. R. Moreira

**Affiliations:** Departamento de Moléstias Infecciosas e Parasitárias, Faculdade de Medicina da Universidade de São Paulo, São Paulo, Brazil; MRC Centre for Global Infectious Disease Analysis, School of Public Health, Imperial College London, London, United Kingdom; Environmental Genomics, Vale Institute of Technology, Belém, PA, Brazil; Departamento de Química Fundamental, Instituto de Química, Universidade de São Paulo, São Paulo, Brazil; Programa Interunidades de Pós-Graduação em Bioinformática, Universidade de São Paulo, São Paulo, Brazil; Faculdade de Medicina Veterinária e Zootecnia, Universidade de São Paulo, São Paulo, Brazil; Laboratório Multiusuário de Doenças Infecciosas e Parasitárias (LADIP) – Universidade Estadual de Montes Claros (Unimontes), Montes Claros, Minas Gerais, Brazil; Laboratório de Comportamento de Insetos, Instituto Federal do Norte de Minas Gerais – campus Salinas, Salinas, Minas Gerais, Brazil; Coordenação-Geral de Vigilância das Arboviroses, Departamento de Doenças Transmissíveis, Secretaria de Vigilância em Saúde e Ambiente, Ministério da Saúde, Brazil; Programa de Pós-graduação em Conservação e Manejo da Vida Silvestre, Unimontes, Montes Claros, Minas Gerais, Brazil; Departamento de Epidemiologia, Faculdade de Saúde Pública da Universidade de São Paulo, São Paulo, Brazil; Laboratório de Mosquitos Transmissores de Hematozoários, Instituto Oswaldo Cruz, Fiocruz, Rio de Janeiro, Brazil; Centro de Excelência de Pesquisa em Saúde – CEPS, Universidade Estadual Montes Claros (UNIMONTES), Montes Claros, MG, Brasil; Instituto Todos pela Saúde, São Paulo, Brazil

**Author notes:** These authors contributed equally to this study (Nuno R. Faria, Filipe R. R. Moreira). **Corresponding author address:** School of Public Health – Imperial College London, Wood Lane, White City, London, W12 0BZ, UK; Instituto Todos pela Saúde, Avenida Paulista, 1938, 01310-942, São Paulo – SP, Brasil.

**Keywords:** Canine distemper virus, neotropical primates, cross-species transmission, host-switching, spillover, hemagglutinin protein, One Health

## Abstract

Canine distemper virus (CDV; *Morbillivirus canis*) is a multi-host paramyxovirus primarily maintained in the order Carnivora. However, recent outbreaks in Neotropical primates suggest recurrent cross-species spillover into non-human primates. Between May and July 2022, seven marmosets (*Callithrix penicillata*) were found dead in urban and peri-urban areas of Montes Claros, Minas Gerais, Brazil. Viral metagenomic sequencing recovered two near complete CDV genomes directly from liver samples. Maximum likelihood and Bayesian phylogenetic analyses placed these genomes within the South American CDV lineage and supported at least three independent zoonotic introductions into *Callithrix* populations. Domestic dogs (*Canis lupus familiaris*) were inferred as the most likely source and the crab-eating fox (*Cerdocyon thous*) as a probable bridge host. Ancestral-state reconstruction identified a substitution at position 519 of the hemagglutinin (H) protein (R519I in the two primate clades and R519G in a singleton primate sequence). *In silico* modelling of the marmoset signaling lymphocytic activation molecule (SLAM) complex was consistent with a modest reorganisation of the receptor-binding head in the 519-substituted variants (ipTM = 0.82), generating a testable hypothesis that this residue may contribute to adaptation at the primate SLAM receptor interface and is associated with altered predicted docking orientation for the receptor. Together, these findings suggest that residue 519 is a candidate marker of CDV adaptation to Neotropical primates and emphasise the importance of integrating genomic surveillance with domestic dog vaccination efforts under a One Health framework to reduce the threat of CDV spillover to vulnerable wildlife populations.

## 1 – Introduction

Canine Distemper Virus (CDV), formally classified as *Morbillivirus canis*, is a highly contagious, multi-host, negative-sense single-stranded 15.4Kb RNA virus that belongs to the genus *Morbillivirus*, family *Paramyxoviridae*^1^. It causes severe multisystemic disease, characterized by respiratory, gastrointestinal, and often fatal neurological symptoms, frequently resulting in immunosuppression and high mortality rates in susceptible species^2^. The CDV genome encodes six structural proteins: nucleocapsid (N), phosphoprotein (P), matrix (M), fusion (F), hemagglutinin (H), and large (L) polymerase^3^. H protein mediates viral attachment to host cellular receptors, Signaling Lymphocytic Activation Molecule (SLAM/CD150) and Nectin-4 (PVRL4)^4–7^, and is widely used for CDV phylogenetic classification ^8–10^.

In Brazil, CDV is believed to be maintained endemically in domestic dog populations, from which viral spillover may occur into wildlife^9,11^. Although historically associated with Carnivora, CDV has increasingly been reported in non-human primates (NHP), particularly Neotropical marmosets of the genus *Callithrix*^9,12,13^. These small-bodied primates often inhabit peri-urban forest fragments where contact with unvaccinated dogs and synanthropic carnivores may facilitate interspecies transmission^14,15^. CDV outbreaks in *Callithrix* populations are typically characterized by severe neurological disease and high mortality, posing a significant threat for wildlife conservation and public health at the domestic animal–wildlife interface^15,16^.

The ability of CDV to jump between mammalian orders is largely dictated by molecular adaptations within the H protein, particularly at sites that interact with the host’s SLAM receptor^9,17^. The receptor-binding domain of H is subject to positive selection, reflecting ongoing optimization of viral attachment to diverse host species^17^. Consistent with this, multiple amino acid substitutions within the receptor-binding interface have been associated with host adaptation in non-canid wildlife, including large felids, mustelids, and non-human primates^9,15–21^. Among these, substitutions at residue 519 have been reported in CDV infecting Neotropical primates and other non-canid hosts, suggesting that this position may contribute to host-specific adaptation, although the available evidence remains limited^9,16,17,22^. Identifying molecular signatures associated with host switching is important for understanding spillover risk and the long-term establishment of CDV in novel hosts.

Here, we describe the genomic characterization of near-complete CDV genomes obtained from *Callithrix penicillata* specimens during a recent epizootic in Brazil. Using viral metagenomic sequencing, we reconstructed near-complete CDV genomes and characterized their genetic diversity and evolutionary relationships. We then applied Bayesian time-scaled analyses to estimate patterns of host switching across the domestic–wildlife interface, and performed structural inferences to assess the potential functional impact of recurring non-synonymous mutations associated with primate clades. Together, these analyses provide new insights into the evolutionary dynamics of CDV host switching and identify residue 519 of the H protein as a candidate molecular determinant of adaptation to Neotropical primates.

## 2 – Material and Methods

### 2.1 – Ethics statement

This study involved the collection and handling of biological samples, which were approved by the Animal Ethics Committee of the Faculty of Medicine of the University of São Paulo under protocol 1849/2022, and by the Authorization and Information System on Biodiversity (Sisbio) under protocol 82076.

### 2.2 – Study population and Specimen Collection

Seven marmosets (*Callithrix penicillata*) were found dead in urban and peri-urban areas of Montes Claros, Minas Gerais, Brazil, by local residents. Necropsies and histopathological examinations were performed to investigate the cause of death. Geographic coordinates of collection sites, sampling dates, and additional metadata are provided in **Supplementary Table S1**.

As necropsy and histopathological findings did not establish a definitive cause of death, viral metagenomic analyses were performed on liver, mesenteric lymph node, and fecal samples from four animals with adequate preservation statuses. The remaining three specimens were excluded due to advanced decomposition and severe autolysis, probably due to the time elapsed between death and sample collection. Selected samples were immediately preserved in RNA*later* (ThermoFischer) and maintained at temperatures between −8 °C and 2 °C for 2 to 7 days. They were then transported on dry ice to the Institute of Tropical Medicine of São Paulo (IMT-SP). Upon arrival, samples were stored at −80 °C in an ultralow-temperature freezer until further processing for viral metagenomic analysis.

### 2.3 – Histopathological examination

Histopathological analyses were performed on formalin-fixed tissue samples from multiple organs, including liver, heart, lung, brain, kidney, and spleen. Following fixation in 10% neutral buffered formalin, tissues were routinely processed, dehydrated through graded alcohols, cleared in xylene, and embedded in paraffin wax. Paraffin blocks were sectioned at 3–5 µm using a microtome, and sections were mounted on glass slides. For histological evaluation, sections were stained with hematoxylin and eosin (H&E) and examined under optical microscopy. Histological assessment was partially limited by marked autolysis observed in several samples, particularly in liver tissue.

Immunohistochemistry (IHC) was performed on liver sections for the detection of yellow fever virus (YFV) antigens, following the protocol previously described by Travassos et al. (1994)^23^. Briefly, sections were subjected to antigen retrieval, followed by blocking of endogenous peroxidase activity, and incubated with a mouse monoclonal anti-YFV primary antibody (produced by the INI). A biotinylated anti-mouse secondary antibody and a streptavidin–biotin detection system were then applied, with chromogenic revelation and hematoxylin counterstaining.

### 2.4 – SMART-9n-ONT Based Viral Metagenomics

Total RNA was extracted from liver and mesenteric lymph nodes using QIAamp RNA Blood Mini Kit (Qiagen), and faecal samples using the QIAamp Viral RNA Mini Kit (Qiagen), following the manufacturers’ protocols with minor modifications. Briefly, tissue fragments (∼3 mm) were homogenized in an RLT buffer (Qiagen) supplemented with 0.1% β-mercaptoethanol using a sterile plastic pestle and incubated at 56 °C for 20 min prior to column-based purification. For faecal samples, approximately 0.5 g of material was diluted in 2 mL of 0.9% NaCl, vortexed, and centrifuged to clarify the suspension; 280 µL of the supernatant was then used for extraction following the viral RNA protocol with AVL buffer. All extracts were subjected to DNase treatment using TURBO™ DNase (Invitrogen™, Thermo Fisher) to remove residual genomic DNA, followed by RNA purification and concentration using the RNA Clean & Concentrator™ kit (Zymo Research), according to the manufacturers’ instructions. Purified RNA was eluted in nuclease-free water and stored at −80 °C until further use.

cDNA synthesis, random amplification, and library preparation for viral metagenomic sequencing were performed following the SMART-9N protocol as described by Morales et al. (2019)^24^. Briefly, 11 µL of extracted nucleic acids were annealed with 1 µL of SMART oligo (2 µM) at 65 °C for 5 min, followed by reverse transcription using SuperScript IV reverse transcriptase in the presence of strand-switching primers (2 µM), RNase inhibitor, dNTPs, and 100 mM dithiothreitol, with incubation at 42 °C for 90 min and enzyme inactivation at 70 °C for 10 min. The resulting cDNA was subjected to random amplification using LongAmp Taq 2× Master Mix with barcoded primers under the following cycling conditions: initial denaturation at 95 °C for 45 s; 26 cycles of 95 °C for 15 s, 56 °C for 15 s, and 65 °C for 5 min; and a final extension at 65 °C for 10 min. Amplified products were purified using AMPure XP beads (1:1 ratio), quantified using the Qubit dsDNA HS Assay, and pooled to obtain an equimolar library (∼400 fM). Rapid adapter ligation was performed at room temperature for 5 min prior to sequencing on an R9.4.1 flow cell (FLO-MIN106D) using a GridION platform (Oxford Nanopore Technologies, ONT). Basecalling and demultiplexing were conducted in real time using Dorado Basecaller (ONT) in high-accuracy mode, with barcode and primer trimming enabled and a minimum Q-score threshold of 9.

All resulting files were submitted to the NCBI Sequence Read Archive (SRA). Sequencing statistics, quality metrics, and the corresponding SRA identification numbers for each sample are provided in **Supplementary Table S2**.

### 2.5 – *In silico* virus identification and viral genome assembly

Raw sequencing reads were processed using a custom pipeline. Briefly, we used minimap2 v2.28-r1209^25^ to map reads against the reference genome of *Callithrix penicillata* (NCBI accession GCA_046862485.1), keeping only unmapped reads for subsequent analyses. Unmapped sequences were submitted to a *de novo* assembly step with MEGA-HIT v1.2.9^26^ and resulting contigs were classified using DIAMOND^27^ with the BLASTx option, and parameters –-ultra-sensitive and e-value 1 x 10^−10^. This analysis used the viral proteins NCBI RefSeq database^28^. Contig sequences preliminarily classified as viruses were subjected to a polishing step with Medaka 2.2.1 (available at: https://github.com/nanoporetech/medaka, last accessed 12 August 2026) and then submitted to a secondary similarity search against a broader database (NCBI nr database^29^). To remove false positive results, we implemented an index-hopping filter, which assumed that a viral detection was only valid if the sample displayed at least 0.5% of the highest read count across all samples in the sequencing run. Complementarily, we also used a negative-control based method to remove false positives, based on the premise that the signal for virus in a sample (aggregated number of reads) should be at least 100x higher than in the negative control.

For the identified viruses, a reference genome assembly was performed with ViralUnity (https://github.com/InstitutoTodosPelaSaude/ViralUnity, last accessed 12 April 2026). Briefly, reads were mapped against the reference viral genome with minimap2 v2.28-r1209^25^ and the resulting bam files were indexed and sorted with samtools v1.23^30^. Variant calling was then performed with Clair 3 v1.2^31^, while variant filtering and consensus sequence inference was performed with bcftools v1.21^30^. Sites with sequencing coverage inferior to 10x were masked. Finally, relevant assembly statistics (number of reads, number of mapped reads, sequencing depth and horizontal coverage) were summarized using custom python scripts included in the ViralUnity pipeline.

### 2.6 – Taxonomic identification of primate species through sequence data analyses

We also leveraged the generated metagenomic data to investigate whether residual host sequencing material could be used to confirm the identity of the morphologically identified primate species, *Callithrix penicillata*. We performed this analysis because the *Callithrix* genus presents morphological overlap in species morphology and a complex landscape of hybrid animals^14^, rendering species-level taxonomic identification uncertain. We therefore inferred the mitochondrial genomes of the four specimens included in this study to perform taxonomic identification through molecular analysis. Briefly, we mapped generated *fastq* files to a *Callithrix jacchus* mitochondrial genome (NCBI accession: NC_025586.1) with minimap2 v2.31^25^, and then inferred a consensus mitochondrial genome sequence with Medaka v2.2.1.

The assembled mitochondrial genomes were then annotated with MitoS v.2.1.10^32^. Per-gene coverage was highly uneven across the assembled draft mitochondrial genomes, with most protein-coding genes showing negligible or no coverage in at least one sample. The genes ATP8, COX1, CYTB, 12S and 16S displayed the most consistent coverage across all samples (100%, 26.3–43.9%, 22.9–43.7%, 80.2–100%, and 95.8–100% of gene length, respectively) and were therefore selected for similarity-search analyses with BLAST^33^, followed by phylogenetic inference. These five gene sequences for each specimen were queried against the nt core NCBI database using BLASTn on the web interface, restricted to family Callitrichidae (taxid: 9480).

A reference phylogenetic dataset was assembled comprising 44 publicly available *Callithrix* mitochondrial genomes retrieved from GenBank (**Supplementary Table 3**), representing all currently recognized species in the genus (*C. jacchus*, *C, penicillata*, *C. kuhlii*, *C. geoffroyi*, and *C. aurita*), together with *Cebuella pygmaea* as outgroup. APT8, COX1, CYTB, 12S, and 16S sequences were extracted from the corresponding reference genomes based on GenBank/RefSeq annotations, combined with sequences from the four specimens, and subsequently aligned using MAFFT v7.505^34^, concatenated, and partitioned by gene. A maximum-likelihood tree was inferred with IQ-Tree2 v2.3.6^35^, using the TPM3u+F+G4 model across all partitions, with node support assessed via 5,000 ultrafast bootstrap replicates and 1,000 SH-aLRT replicates. To formally test alternative species-assignment hypotheses, we performed the Approximately Unbiased (AU) topology test^36^, comparing the unconstrained maximum-likelihood topology against constrained topologies enforcing reciprocal monophyly of each specimen with each candidate congener species. The AU tests were implemented with 10,000 RELL bootstrap replicates^37^.

### 2.7 – Phylogenetic contextualization of novel CDV sequences

To contextualize the new CDV genomes, we collated a comprehensive reference dataset encompassing all near-complete genomes available on NCBI Genbank as of April 1st, 2026. We refer to near-complete genomes as all CDV sequences with at least 15,000 bp. Only sequences with collection dates and host information available were kept in the dataset. New and reference sequences were aligned with MAFFT v7.526^34^ and 5’ and 3’ UTR regions were manually trimmed. A maximum-likelihood phylogenetic inference was performed with IQ-Tree v3.0.1^35^ under the GTR+F+G4^38^ nucleotide substitution model, as selected by Model-Finder^39^. Ultra-fast bootstrap was used to assess branch statistical support^40^. Motivated by the results from this analysis, we performed a secondary phylogenetic inference focusing specifically on a clade comprehending the Europe/South America 1 genotype. This second dataset had its temporal signal verified through a root-to-tip regression with TempEst v1.5.3^41^.

### 2.8 – Bayesian time-scaled phylogenetic reconstruction

To further investigate evolutionary patterns underlying the CDV outbreaks in *Callithrix*, we performed a time-scaled phylogenetic inference using BEAST X^42^ and the BEAGLE library V4.0.1^43^ for accelerated likelihood computation. This analysis used a GTR+F+G4 nucleotide substitution model^38^, a strict molecular clock model with a skygrid tree prior under the recently implemented Hamiltonian Monte-Carlo operator^44,45^. We jointly performed a discrete two-trait analysis (DTA) to reconstruct the dynamics of host-switching and geographic dispersion patterns from this dataset. The host-switching analyses were performed under two parameterizations: one with two states, reflecting host order (Carnivora and Primates – DTA host model 1) (**Figure 3**), and another with 9 states, reflecting host genus (*Callithrix*, *Canis*, *Cerdocyon*, *Lutra*, *Martes*, *Mustela*, *Procyon*, *Pusa* and *Vulpes –* DTA host model 2) (**Supplementary Figure S1**). Likewise, the phylogeographic reconstruction used distinct parameterizations, one with three states, reflecting continents (South America, Africa and Europe – DTA spatial model 1; shown in **Supplementary Figure S2 panel A**), and another with seven states based on countries (Argentina, Brazil, Gabon, Germany, Hungary, Italy and Uruguay – host model 2, **Supplementary Figure S2 panel B**). The number of sequences in each category for each parameterization is summarized in **Supplementary Table S4**.

Each analysis was performed on duplicate Markov Chain Monte Carlo runs with 100 million generations, sampling every 10,000 steps. Mixing and convergence were verified on Tracer v1.7.2^46^. Logs and trees were combined using LogCombiner v10.5.0 and a maximum-clade credibility tree was inferred with TreeAnnotator v1.10.4. Trees were visualized using the PearTree software v1.3.1 (available at: https://github.com/artic-network/peartree, last accessed 17 August 2026).

### 2.9 – Ancestral states reconstruction and selection analyses

To further investigate patterns of molecular evolution underlying host-switching in CDV from carnivores to primates, we performed phylogenetic ancestral state reconstructions on amino acid sequences from the H protein using TreeTime^47^. We also used codon-evolution models to investigate signs of positive selection with HyPhy^48^. Specifically, we used the Mixed-Effects Evolutionary Model (MEME)^49^ and the Fixed-Effects Likelihood (FEL) model^50^ to identify sites displaying signs of adaptive evolution. These models were used to analyze both the full dataset and the dataset corresponding to the subtree encompassing the clade of interest. A significance level of 0.01 was determined for these analyses.

### 2.10 – Structural analyses of the CDV Hemagglutinin protein

To investigate the structural basis of host adaptation, we performed *in silico* protein–protein docking and interaction analysis using AlphaFold3^51^. The amino acid sequence of the Canine distemper virus (CDV) Hemagglutinin (H) protein ectodomain (residues 59–607) was modeled in complex with the V-domain of the marmoset (*Callithrix jacchus*, accession number: XP_002760222.4) SLAM (CD150) receptor (residues 28–140), as this domain is the primary determinant of *Morbillivirus* host specificity^5,52–54^. No SLAM annotation for the *C. penicillata* reference genome existed at NCBI. We then conducted three separate molecular dockings to compare the structural properties of the ancestral carnivore-associated strain (carrying the 519R wild-type residue; NCBI accession number MH426739) and the primate-associated strains (carrying the 519I/G mutations, accession numbers PZ407655 and PQ464015, respectively). Structural confidence was assessed using the interface Predicted Template Modeling (ipTM) score and Predicted Aligned Error (PAE) plots to quantify the reliability of the docked complex^55^. PAE matrices were extracted from the AlphaFold3 output and plotted as heatmaps using a custom Python script (Matplotlib v3.10). The resulting models were visualized and analyzed in UCSF ChimeraX v1.9^56^ to map the physical distance between the mutation site and the host receptor and to identify specific residues at the binding interface within a 4.0 Å radius.

## 3 – Results

### 3.1 – Epizootic description

Seven dead non-human primates identified as *Callithrix penicillata* were found by local residents in urban and periurban areas of the municipality of Montes Claros between May and July 2022. The carcasses were collected by the regional Zoonosis Control Center (*Centro de Controle de Zoonoses – CCZ*) and sent to National Institute of Infectology Evandro Chagas, where necropsies and histopathological examinations were performed to investigate the cause of death. No relevant histopathological alterations were observed in the evaluated tissues. In addition, considering the epidemiological importance of yellow fever virus (YFV) in non-human primate populations in Brazil, all tested samples were assessed for YFV antigen by immunohistochemistry (IHC). All samples were negative, excluding YFV infection as the cause of death.

### 3.2 – Portable metagenomic sequencing provides evidence implicating CDV in mortality events affecting *Callithrix penicillata* in Minas Gerais

To identify the aetiology of the reported potential epizootic in marmosets, we performed viral metagenomic sequencing using the SMART-9N protocol with the GridIon sequencer. Due to the advanced state of decomposition of three carcasses, only four animals were considered suitable for investigation. We sequenced a total of 12 samples (faeces, mesenteric lymph nodes, livers), from 4 animals, and generated a total of 6 million reads, with an average throughput per sample 516,496.70 reads (standard deviation: 701,638.7; range: 2,798 – 2,245,895) (**Table 1**). These sequences were submitted to a comprehensive metagenomic analysis pipeline, which performed host depletion, *de novo* assembly and two rounds of similarity search against distinct databases, also applying strict filters to remove false positive detections. As a result, we identified two viral families across all samples, including *Paramyxoviridae* (CDV), in two liver samples, and *Iflaviviridae* (Sanya iflavirus), in one fecal sample. Notably, CDV was identified in liver samples from two animals collected in the Sapucaia region on 2022-05-23 and 2022-05-27. We then assembled two near-complete CDV genomes (99%> horizontal coverage), which are available at GenBank under accession numbers PZ407655 and PZ407656. Complete genome assembly statistics are available in **Supplementary Table S5**.

**Table 1.** Sequencing summary metrics for metagenomic samples obtained from faeces, lymph, and liver tissues. Values are presented as median [Q1–Q3] for each sample type (n = 4 samples per tissue). Sequencing metrics included total number of reads, median read length (bp), N50 (bp), and median quality score. Q1 and Q3 represent the first and third quartiles, respectively, corresponding to the interquartile range (IQR). The number of reads is expressed in thousands (K).

| Sample type | Liver | Lymph | Faeces |
| --- | --- | --- | --- |
| **Number of reads\*** | 611,879<br>[389,390–931,581] | 124,505<br>[116,675.3–170,845.5] | 196,391<br>[140,920–732,853.3] |
| **Read length (bp)\*** | 289.5 [256.3–324.5] | 216.5 [214.8–218.8] | 278.5 [270.5–292.5] |
| **N50 (bp)\*** | 358 [342.3–373.5] | 247.5 [238.3–262] | 306.5 [293–327.8] |
| **Median quality score\*** | 12.9 [12.5–13.1] | 8.8 [8.7–9] | 8.6 [8.6–8.9] |

In addition to viral identification, sequence data analysis allowed the reconstruction of partial mitochondrial genomes across all specimens, with an average coverage breadth of 45.5% (**Table 2**; **Figure 1A**). Similarity searches (BLASTn) using both individual genes and concatenated gene sequences (ATP8+COX1+CYTB+12S+16S) mostly retrieved *C. penicillata* as the top hit (**Table 2**). We also submitted sequences to maximum-likelihood phylogenetic reconstructions. Inference using the concatenated (ATP8+COX1+CYTB+12S+16S) alignment placed all four specimens within a single, well-supported clade nested inside *Callithrix penicillata* (**Figure 1B**). Single-gene analyses showed partial and, in some cases, conflicting support for this placement (**Supplementary Figure S3**). To evaluate phylogenetic uncertainty, AU topology tests were used to formally evaluate alternative species assignments on the concatenated alignment, significantly rejecting monophyly with *C. jacchus* (ΔlnL = 342.7, p < 0.0001) or *C. kuhlii* (ΔlnL = 75.9, p < 0.0001) and supporting *C. penicillata* as the best-supported maternal lineage (**Table 2**). Together, the concatenated phylogenetic analysis, AU topology tests, and the majority of BLASTn analyses consistently support taxonomic identification of *C. penicillata*, confirming morphological classification (**Table 2**, **Figure 1**).

**Figure 1.**
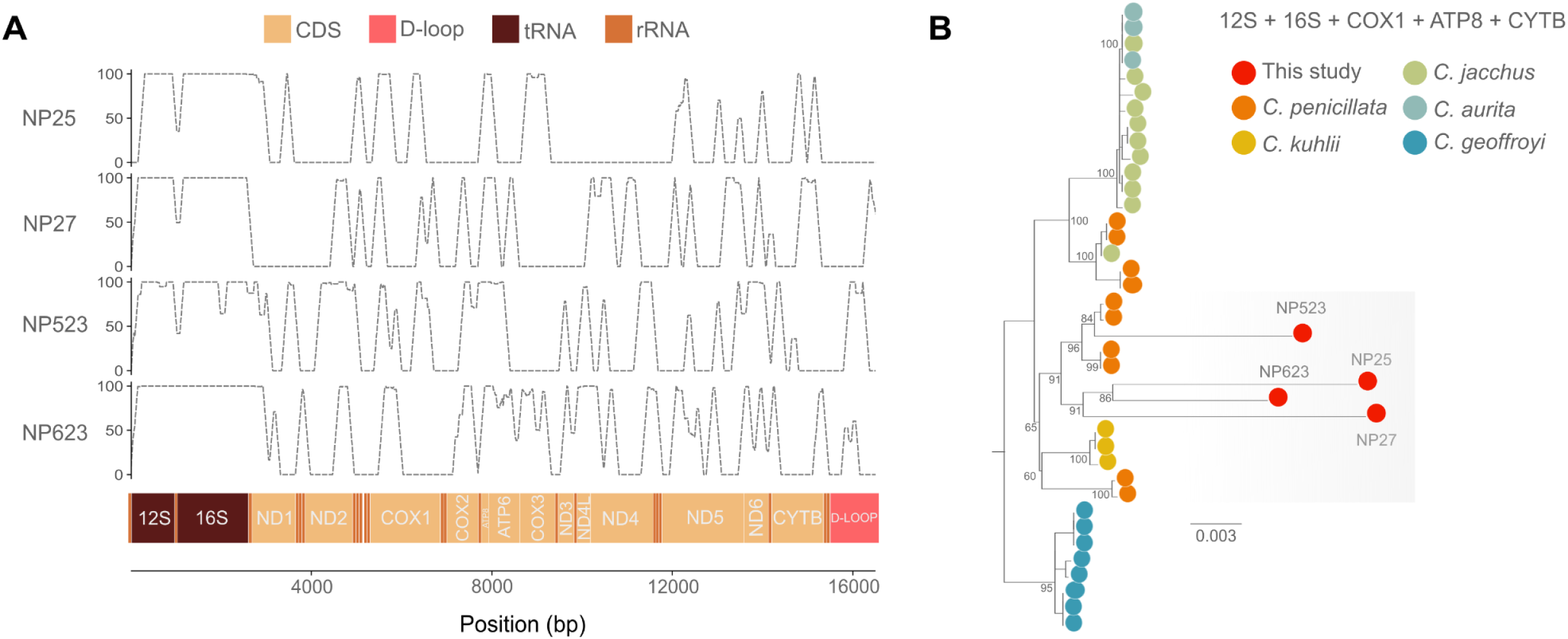
Recovery of host mitochondrial genomes from metagenomic sequencing and phylogenetic placement of the study samples. **(A)** Sequencing depth across the mitochondrial genome obtained by mapping metagenomic sequencing reads from each sample to the *Callithrix jacchus* reference mitochondrial genome. The mitochondrial genome organization is shown below the coverage plots, including protein-coding genes (CDS), rRNAs, tRNAs, and the D-loop/control region. **(B)** Maximum likelihood phylogenetic tree inferred from partial mitochondrial genome sequences recovered in this study together with representative *Callithrix* mitochondrial genomes. This dataset corresponds to the concatenation of individual gene alignments (12S+16S+COX1+ATP8+CYTB). Branches are in the scale of substitutions per site. Ultrafast bootstrap values are exhibited along the most relevant nodes of the tree. Tip shapes are colored according to the record species or refer to samples generated in this study. The visualization corresponds to a subtree, not including a distantly related *C. aurita* clade. The new sequences (red circles; NP25, NP27, NP523, and NP623) cluster with *C. penicillata*, supporting their assignment to this species.

**Table 2.** Per-sample mitochondrial genome and pergene coverage, BLASTn similarity search result, phylogenetic placement and approximately unbiased (AU) topology test.

| Sample | mt-DNA cov. (%) | Gene cov. (%)<br>ATP8 / COX1 /<br>CYTB /<br>12S / 16S | BLASTN top hit<br>Concatenated /<br>COX1 / CYTB / 12S /<br>16S (identity %) | Phylogenetic<br>placement<br>(all genes) | Phylogenetic AU test<br>support vs.<br>alternative |
| --- | --- | --- | --- | --- | --- |
| NP25 | 33.1 | 100.0 / 43.9 /<br>32.1 /<br>80.2 / 99.8 | *C. penicillata* (99.5%)<br>*C. penicillata* (96.2%)<br>*C. penicillata* (98.5%)<br>*C. penicillata* (99.2%)<br>*C. penicillata* (99.6%) | *C. penicillata* | p = 1 (*penicillata*);<br>rejected: *jacchus*,<br>*kuhlii* (p < 0.0001) |
| NP27 | 40.4 | 100.0 / 32.1 /<br>43.7 /<br>99.0 / 99.9 | *C. penicillata* (99.32%)<br>*C. penicillata*/*kuhlii*,<br>tied (100%)<br>*C. penicillata* (93.4%)<br>uninformative (99.2%)<br>*C. penicillata* (99.3%) | *C. penicillata* | p = 1 (*penicillata*);<br>rejected: *jacchus*,<br>*kuhlii* (p < 0.0001) |
| NP523 | 47.9 | 100.0 / 40.5 /<br>22.9 /<br>95.0 / 95.8 | *C. penicillata* (99.2%)<br>*C. penicillata* (99.6%)<br>*C. penicillata*/*jacchus*,<br>tied (97.1%)<br>uninformative (95.6%)<br>uninformative(99.2%) | *C. penicillata* | p = 1 (*penicillata*);<br>rejected: *jacchus*,<br>*kuhlii* (p < 0.0001) |
| NP623 | 50.7 | 100.0 / 26.3 /<br>23.7 /<br>100.0 / 100.0 | *C. penicillata* (99.7%)<br>*C. penicillata* (92.5%)<br>uninformative, tied<br>(96.6%)<br>uninformative (98.8%)<br>*C. penicillata* (99.7%) | *C. penicillata* | p = 1 (*penicillata*);<br>rejected: *jacchus*,<br>*kuhlii* (p < 0.0001) |

### 3.3 – Phylogenetic methods support the occurrence of multiple zoonotic transmissions of CDV to marmosets

Our phylogenetic reconstruction largely reflects previous estimates of the evolutionary history of CDV, with main genotypes and relationships among them reconstructed with high statistical support (UFBoot >= 90) (**Figure 2A**). Our newly generated genomes cluster within the diversity of the Europe/South America 1 genotype, consistent with previously generated genomes characterized from *Callithrix* sp. samples (**Figure 2B**). Notwithstanding, our sequences do not cluster with sequences derived from other primate outbreaks in Brazil, but with a virus characterized from a crab-eating fox (*Cerdocyon* sp.) individual sampled in Brazil in 2014 (accession number MH426739, UFBoot = 68). To further investigate the origins of this epizootic, we first assessed the temporal signal of the subtree encompassing the Europe/South America 1 genotype. Our root-to-tip regression suggested a substantial temporal signal (correlation coefficient = 0.83, R-squared = 0.68, slope = 5.1 x 10^−4^), with no detectable outliers (**Figure 2C**).

**Figure 2.**
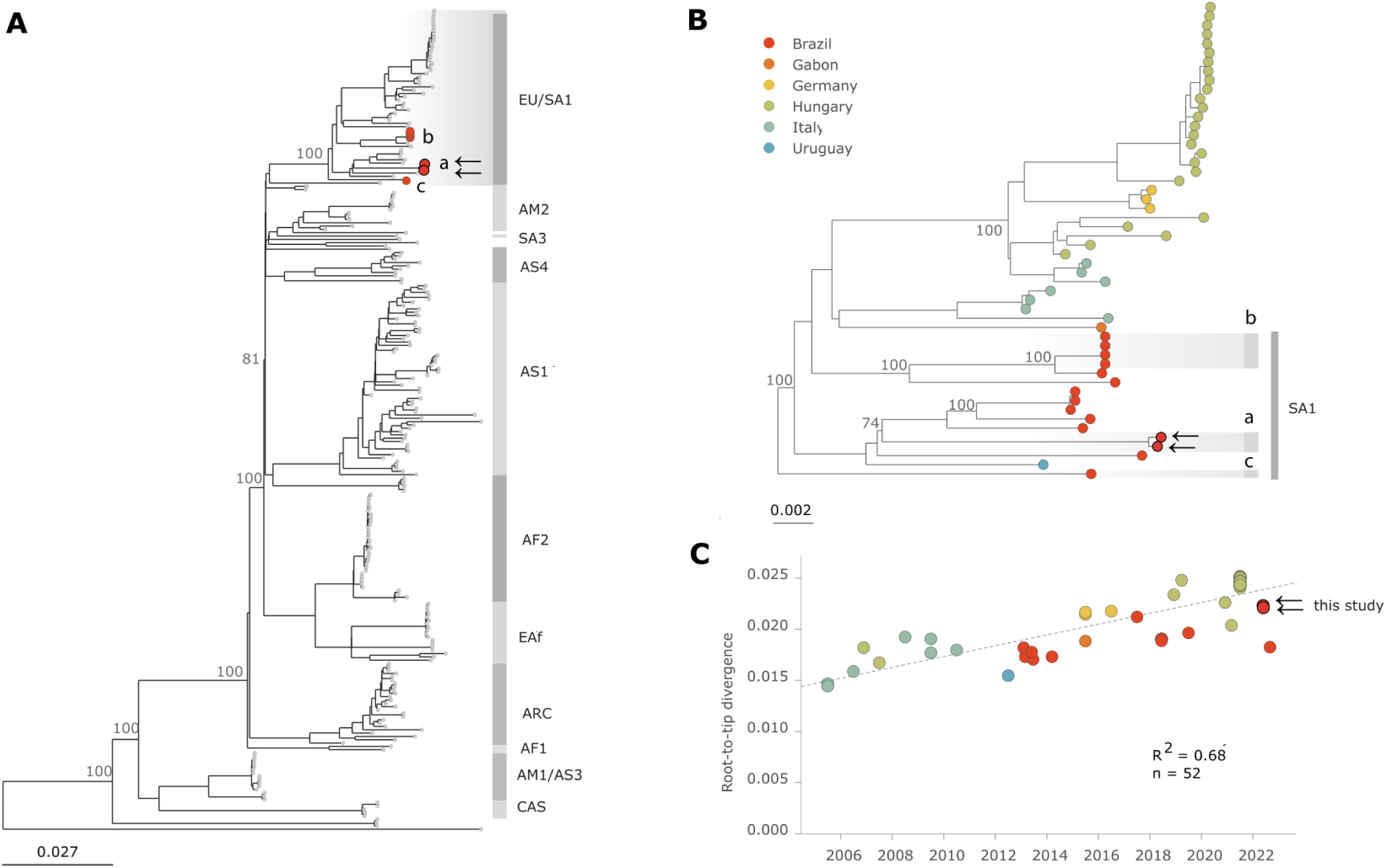
Phylogenetic reconstruction and temporal signal analysis of canine distemper virus (CDV). **(A)** Maximum-likelihood phylogeny inferred from all publicly available complete CDV genomes (>15 kb). Major CDV genotypes are indicated, with sequences obtained from *Callithrix penicillata* highlighted in light red and the genomes generated in this study highlighted in dark red, all of them from Brazil. Ultrafast bootstrap (UFBoot) support values are shown for major nodes. Genotype abbreviations are as follows: EU, Europe; SA, South America; AM, America; AS, Asia; AF, Africa; EAf, East Africa; ARC, Arctic-like; CAS, Caspian. The arrows indicate the samples provenient from this study. **(B)** Geographic distribution of sequences belonging to the Europe/South America 1 genotype. **(C)** Root-to-tip regression analysis of the Europe/South America 1 genotype showing the temporal signal.

Next, we estimated a Bayesian time-calibrated tree along a two discrete trait model considering both host and geography. Our analyses dated the common ancestor of CDV in Europe/South America 1 lineage to around 1933 (95% highest posterior density [HPD]: 21 April 1921 – 14 August 1944) (**Figure 3**). The virus evolutionary rate was estimated around 3.82 x 10^−4^ substitutions per site per year (95% HPD: 3.29 x 10^−4^ – 4.36 x 10^−4^). Both parameterizations (continent and country-based spatial models 1 and 2, respectively) support the ancestor of the whole tree (root) to have existed in South America at this time (spatial posterior probability [pp] > 0.99), later spreading to Europe (pp = 0.88) around 24 October 1967 (95% HPD: 04 January 1964, 01 January 1977), via Italy (pp = 0.36) (**Supplementary Figure S2**). In terms of spread within South America, our country-based analysis supports that all nodes in the region had Brazil as the most likely state, with the country contributing to introductions in Uruguay and Argentina. Because sampling was uneven across locations, we interpret our phylogeographic results as a signal of broad-scale movement patterns rather than as a precise estimate of transition rates.

Our host-switching analysis, which considers hosts as discrete traits, supports that at least three independent introductions of CDV in *Callithrix* populations occurred in Brazil (**Figure 3, Supplementary Figure S2**). Two of these introductions were related to clades from distinct primate epizootics^16^. Clade B is associated with four sequences sampled in 2018 from *Callithrix* sp. individuals in Brasília (Centre-West region), being dated around 26 September 2017 (95% HPD: 22 January 2018 – 06 June 2018, phylogenetic pp = 1, host model 1 pp = 1, host model 2 pp = 1). This clade is most closely related to a virus sampled from a *Cerdocyon thous* individual in 2018 (accession number: PP847349, phylogenetic pp = 0.52), and the tMRCA for the whole group was dated to around 21 October 2009 (95% HPD: 07 August 2008 – 11 May 2012).

**Figure 3.**
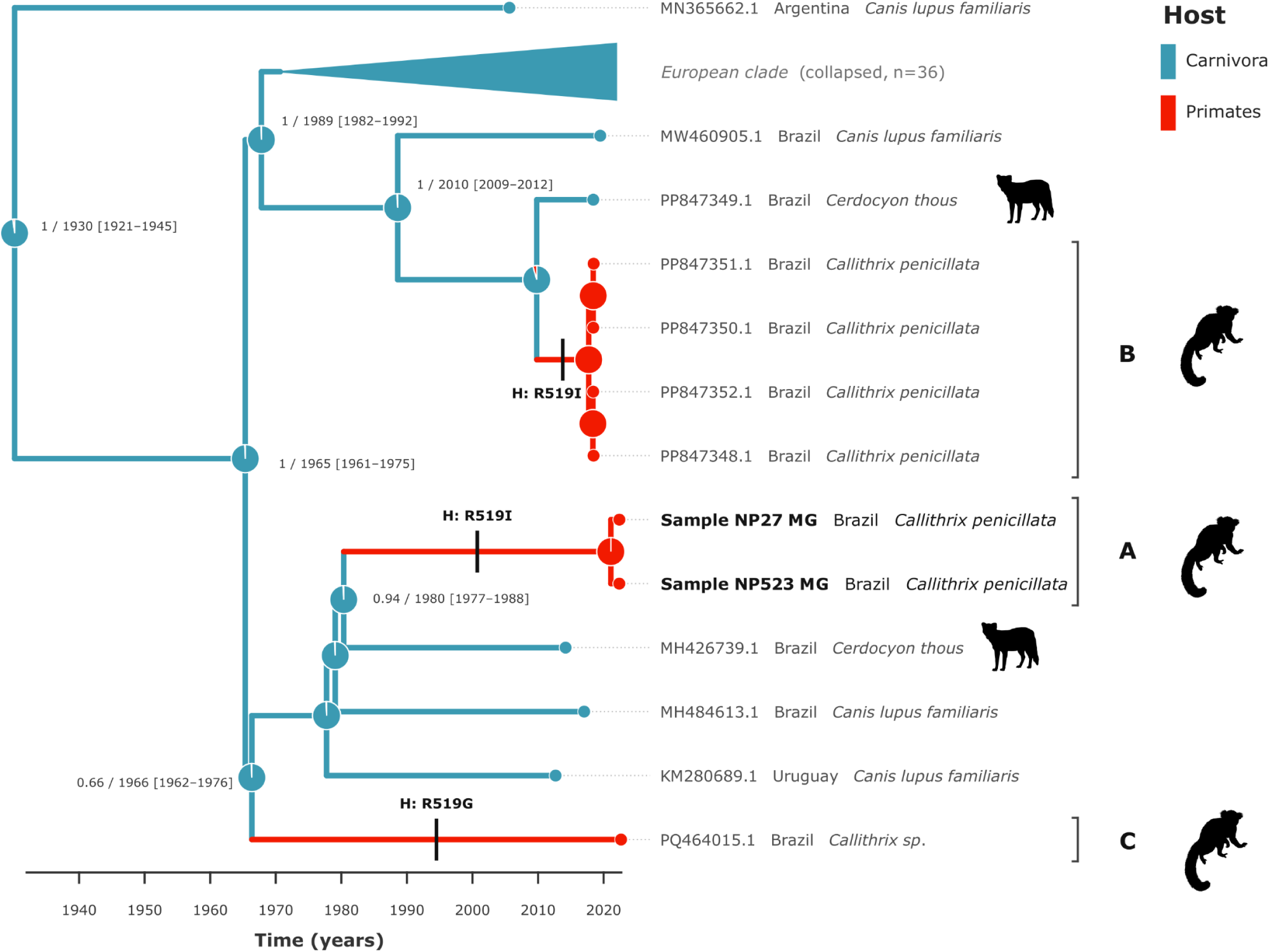
Bayesian time-scaled phylogenetic reconstruction of Canine Distemper Virus (CDV) Europe/South American 1 genotype highlighting zoonotic transitions to primates. The maximum clade credibility tree illustrates the evolutionary relationships of CDV lineages across diverse hosts, with branches color-coded by reconstructed ancestral host states: Carnivora (blue) and Primates (red). Node shapes express uncertainty in ancestral states reconstruction. Branches are scaled to time (years). The European clade was pruned for clarity. Nodes of interest are annotated with posterior probability (PP) values and estimated divergence dates (mean and 95% highest posterior density [HPD] intervals). Transverse dashes indicate the independent occurrence of the R519I/G mutation, a substitution associated with viral adaptation to the primate SLAM receptor. Tip labels include GenBank accession numbers, geographic origin and host species. The topology demonstrates multiple independent spillover events into South American primates (e.g., *Callithrix penicillata*), originating from sympatric carnivore populations including domestic dogs (*Canis lupus familiaris*) and crab-eating foxes (*Cerdocyon thous*). These spillover and downstream transmission events led to the emergence of clades A, B and C, marked on the tree.

The second primate epizootic clade is composed of sequences generated in this study (clade A), sampled in Minas Gerais (Southeast region) in 2022, and was dated to around 08 February 2021 (95% HPD: 08 May 2020 – 11 August 2021, phylogenetic pp = 1, host model 1 pp = 1, host model 2 pp = 1). This clade is related to a virus characterized from a *Cerdocyon thous* individual (accession number: MH426739, pp = 0.73), but the tMRCA for the whole clade is much older (mean: 12 May 1980, 95% HPD: 10 June 1977 – 22 January 1988). The third reconstructed introduction of CDV into *Callithrix* populations is represented by a single sequence, herein named clade C, which diverged from the closest South American clade around 1966 (95% HPD: 11 January 1962 – 30 October 1975).

### 3.4 Recurrent emergence of mutations in the 519 residue of the Hemagglutinin protein

We next investigated whether transitions between carnivore and primate hosts were associated with recurrent genomic signatures. To do this, we restricted our analysis to the Hemagglutinin (H) gene, which encodes the key protein responsible for virus attachment and tropism for morbilliviruses. First, we performed an ancestral state reconstruction and verified whether any amino-acid residue consistently emerged along any of the three branches marking transitions between carnivore and primate hosts. We observed several substitutions that emerged independently along these branches (**Supplementary Figure S4**), some of those with potential biological significance due to their location (ectodomain, sites 59-607). However, only one residue, H-519, exhibited recurrent substitutions across all independently derived primate-associated clades (**Figure 3**). In the two primate clades, the substitution R519I was present, while the singleton sequence displayed the mutation R519G. While other mutations of potential significance were identified, including G530R/S, no other was consistently recovered along all branches leading to clades A, B and C. Trees annotated with the complete set of reconstructed mutations are available on a dedicated GitHub repository (https://github.com/MayaraBertanhe/CDV-Sapucaia, last accessed on 07 Aug 2026).

We then investigated the H gene alignment with selection models (MEME and FEL), and found evidence of natural selection at the molecular level across a number of sites (see **Supplementary Table S6**). On the full dataset, MEME identified three sites with consistent signs of episodic positive selection (p < 0.01), including G104T, L105S/V/Y and T291A/I/M, while none was identified in the smaller dataset (Europe/South America 1 genotype). Similarly, the FEL model identified a single site with consistent signal of positive selection on the full dataset, Y549H, while none was identified in the smaller dataset. The FEL model also identified signals of sites under purifying natural selection on both the full and smaller datasets, comprehending 142 and 10 sites, respectively. None of the mutation signals of positive selection were associated with branches leading to primate clades or sequences.

### 3.5 – Hemagglutinin R519I/G mutations and structural properties of the H-SLAM interface in marmosets

Structural modelling indicated higher-confidence predicted H-SLAM complexes for the R519I/G variants relative to the ancestral R519 background. Specifically, the primate-associated viral H-proteins exhibited higher ipTM score (519I: 0.82, 519G: 0.86) compared to the ancestral carnivore-associated H-protein (0.75) (**Figure 4**). Analysis of the Predicted Aligned Error (PAE) plots corroborated this finding, showing a substantial reduction in positional uncertainty across the virus-receptor interface in the mutant models (**Supplementary Figure S5**). The R519 residue is situated on the lateral surface of the H-protein head at the interface with the SLAM V-domain (**Figure 4A**). Structural visualization revealed that R519 participates in intramolecular hydrogen bonds with neighboring H-protein residues, and mutations at this position substantially alter these local interactions (**Figure 4B**). In the wild-type complex, R519 maintains hydrogen bonds with adjacent H-protein residues (**Figure 4B**, left panel). The R519I and R519G mutations, however, exhibit distinct hydrogen bonding patterns within the H-protein due to the chemical properties of isoleucine and glycine (**Figure 4B**, middle and right panels). These alterations in intramolecular interactions may propagate to affect local protein organization and potentially influence the geometrical configuration of the H-protein at the SLAM V-domain interface, as suggested by the higher ipTM scores in the mutants. However, these observations represent an initial structural hypothesis. The functional significance of these local rearrangements and their impact on viral-receptor interactions requires further validation through molecular dynamics simulations and biochemical assays.

**Figure 4.**
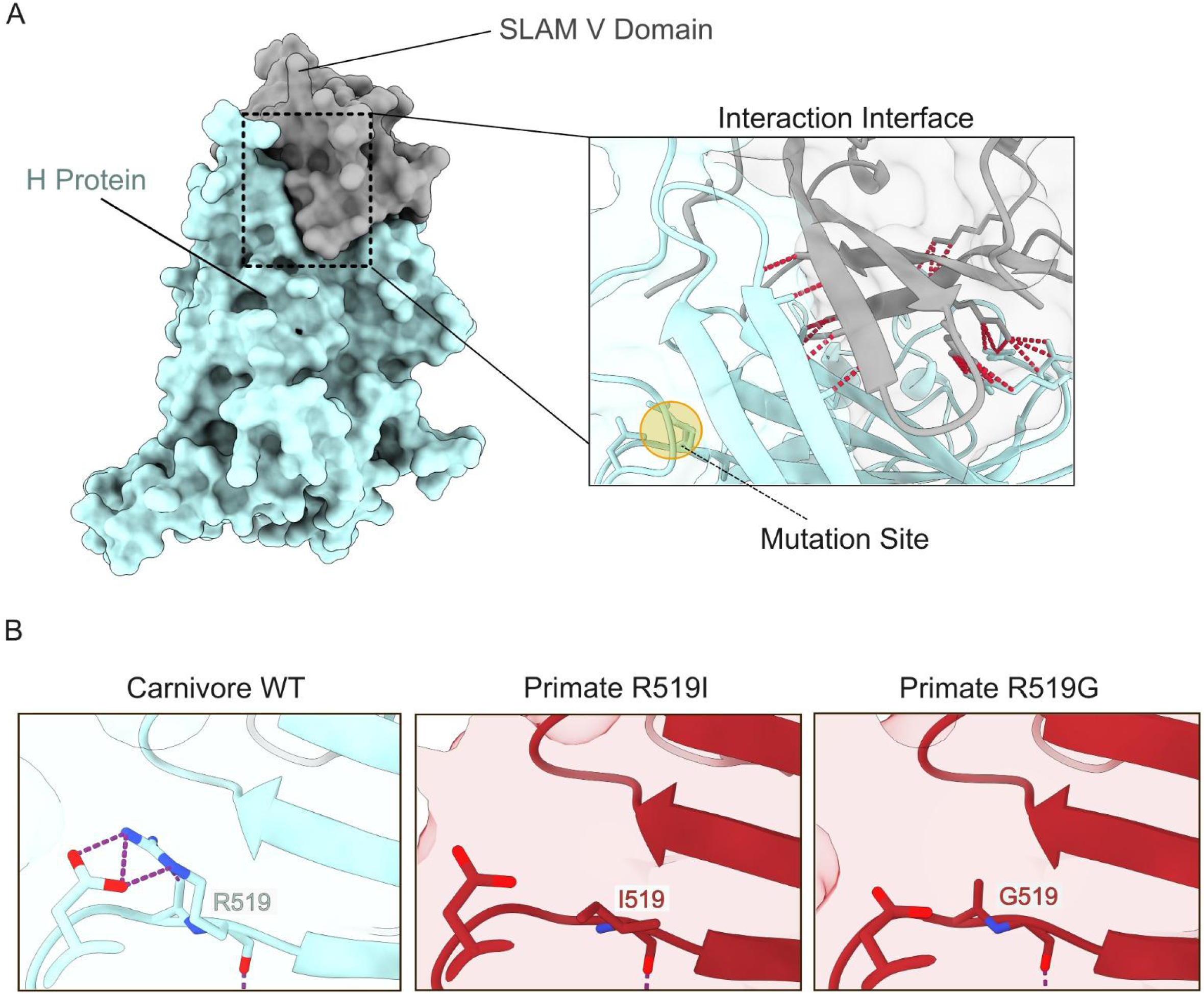
Local Structural Rearrangements in CDV H-protein R519 Variants. **(A)** Overview of carnivore wild-type H-SLAM complex showing the H-protein (blue surface) and SLAM V-domain (gray). The dashed box indicates the region magnified on the right, displaying intramolecular hydrogen bond patterns (cyan dashed lines) within the H-protein. The residue 519 position where mutations are observed is highlighted in yellow. **(B)** Comparative structural detail of the three variants: carnivore wild-type (WT) with R519, primate R519I mutant, and primate R519G mutant. Purple dashed lines represent intramolecular hydrogen bonds within H-protein.

## 4 – Discussion

The identification of Canine Distemper Virus (CDV) as the primary driver of a potential epizootics in *Callithrix penicillata* highlights an expanding threat to Neotropical primate conservation^16^. Using viral metagenomic sequencing, we successfully reconstructed near-complete genomes directly from clinical samples, confirming the high susceptibility of marmosets to local CDV diversity (Europe/South America 1 genotype). Notably, the Brazilian landscape is characterized by the complex co-circulation of multiple wild-type lineages, including CDV South America (SA)-1, SA-2, SA-3, and SA-4^9^. Our findings further reveal a dynamic pattern of recurrent host switching and convergent molecular evolution, particularly involving residue 519 of the hemagglutinin (H) protein, highlighting its potential association with adaptation of CDV to alternative host species.

Our phylogenetic analysis demonstrates that CDV was introduced into *Callithrix* populations through multiple independent zoonotic events rather than a single sustained transmission chain. The close relationship between primate-derived sequences and those from the crab-eating fox (*Cerdocyon thous*) suggests these synanthropic meso-carnivores may act as critical bridge hosts between domestic dog reservoirs and sensitive NHP populations^9,57^. Furthermore, the genetic distance between the 2022 Minas Gerais sequences and their closest available wildlife-derived genomes suggests that substantial unsampled viral diversity persists in wildlife reservoirs. This silent circulation presents a major obstacle to surveillance and emphasizes the need for a One Health multipathogen and viral-agnostic approaches that integrate monitoring of domestic, wild, and primate populations^16^. Additionally, the close temporal, spatial and genetic proximity of the two CDV-infected marmosets, sampled in this study from the same area only four days apart, raises the possibility of marmoset-to-marmoset transmission, a hypothesis previously proposed for CDV in non-human primates^16,58^. However, these associations alone do not constitute direct evidence of transmission between marmosets, and repeated independent spillover events cannot be excluded.

Our data suggests that residue 519 of the H protein represents a candidate marker associated with host-specific adaptation. While residues 530 and 549 are established markers for CDV expansion into non-canid hosts^9,17,18^, Old World primates (Catarrhini) often retain the canid-typical signatures at these sites^9^. Instead, our reconstruction identifies site 519 as a potential adaptive marker for CDV transmission in Neotropical primates, specifically through the R519I substitution^15^. Although standard selection models were unable to detect episodic or pervasive positive selection, this may reflect limited power after rapid fixation following a hard sweep. The consistent presence of these substitutions across all primate branches nevertheless supports their likely biological relevance, although their phenotypic consequences remain to be experimentally determined.

To explore a potential mechanistic basis for this evolutionary pattern, we modelled the interaction between the H protein and the marmoset SLAM receptor. This analysis indicated that substitutions R519I and R519G were associated with more confident H-SLAM complex predictions, reflected by higher ipTM scores (0.82 and 0.86, respectively) than the ancestral variant (0.75). Inspection of the predicted complexes further revealed that these substitutions alter the local hydrogen-bonding network surrounding residue 519, suggesting that they may induce subtle structural rearrangements capable of reorganizing the H-protein interface with the SLAM V-domain. Although ipTM and PAE scores do not directly quantify binding affinity, the observed structural changes are consistent with the hypothesis that substitutions at residue 519 may influence receptor engagement. This interpretation is supported by previous experimental work in lions, in which CDV strains carrying the adapted H-protein genotype (519I/549H) displayed enhanced viral entry into cells expressing lion SLAM compared with viruses retaining the ancestral 519R/549Y residues^17,22,59^. Future studies integrating molecular dynamics simulations, binding free-energy calculations (e.g., MM-PBSA), and functional viral entry assays will be important to determine whether the structural differences observed here translate into measurable changes in receptor affinity and infectivity.

Preventing future CDV spillover events in primates will require robust One Health firewall strategies. Vaccination of domestic dogs remains the principal strategy currently available for reducing CDV circulation and limiting opportunities for cross-species transmission^59^. However, vaccination coverage is frequently insufficient in many regions of Brazil, particularly at peri-urban and rural interfaces where domestic animals and wildlife co-occur^11^. Moreover, the extent to which domestic dog vaccination effectively protects wildlife populations remains poorly understood, especially given the growing evidence that wild carnivores may play an important role in maintaining and disseminating CDV independently of domestic reservoirs, and the fact that these wildlife hosts are not subject to vaccination programs^59^. Given that, surveillance efforts should be strengthened by incorporating CDV testing into routine diagnostic screening of wild animals found dead, whenever feasible.

The ecological context of the Sapucaia region highlights these challenges. *Parque Municipal da* Sapucaia, located in the Serra do Mel region in southern Montes Claros, comprises a forest reserve of more than 300,000 m² embedded within an urbanized landscape, creating extensive opportunities for contact among wildlife, domestic animals, and humans. Such interfaces may facilitate pathogen exchange among species and increase the likelihood of spillover into highly susceptible primate populations. Notably, this scenario parallels observations during yellow fever virus outbreaks in Brazil, where transmission has been concentrated at forest fragments and where genomic surveillance of infected primates and vectors has provided critical insights into viral dispersal pathways and the ecological routes through which pathogens reach susceptible Neotropical primate populations^60^. Together, these findings reinforce the importance of integrated surveillance programmes that combine wildlife monitoring, pathogen genomics, and domestic animal health measures. Although vaccination of domestic dogs alone is unlikely to eliminate spillover risk, it remains the most practical intervention currently available to reduce opportunities for cross-species transmission. Future studies using in vitro entry and binding assays will be essential to experimentally quantify the contribution of the R519I/G substitution to host-specific entry and to further delineate the molecular barriers to morbillivirus emergence in South American primates.

## 5 – Acknowledgements

We thank the *Centro de Controle de Zoonoses de Montes Claros* for allowing us to follow the necropsies of the specimens and for facilitating the collection and acquisition of biological samples used in this study. We also thank the collaborators from the *Laboratório Multiusuário em Doenças Infecciosas e Parasitárias* (LADIP), *Universidade Estadual de Montes Claros* (Unimontes), for their valuable assistance and support throughout the study.

## 8 – Data availability

Raw sequencing reads generated on this study are publicly available on the NCBI short-read archive (SRA) under accessions SRR38833314-SRR38833323 (Bioproject PRJNA1467027, Biosamples SAMN60298444-SAMN60298453). Consensus genome sequences are available on NCBI GenBank under accessions PZ407655 and PZ407656. Alignments, trees, beast XMLs and log files are available on a dedicated GitHub repository: https://github.com/MayaraBertanhe/CDV-Sapucaia (last accessed on 07 Aug 2026).

---

**Supplementary Figure S1.**
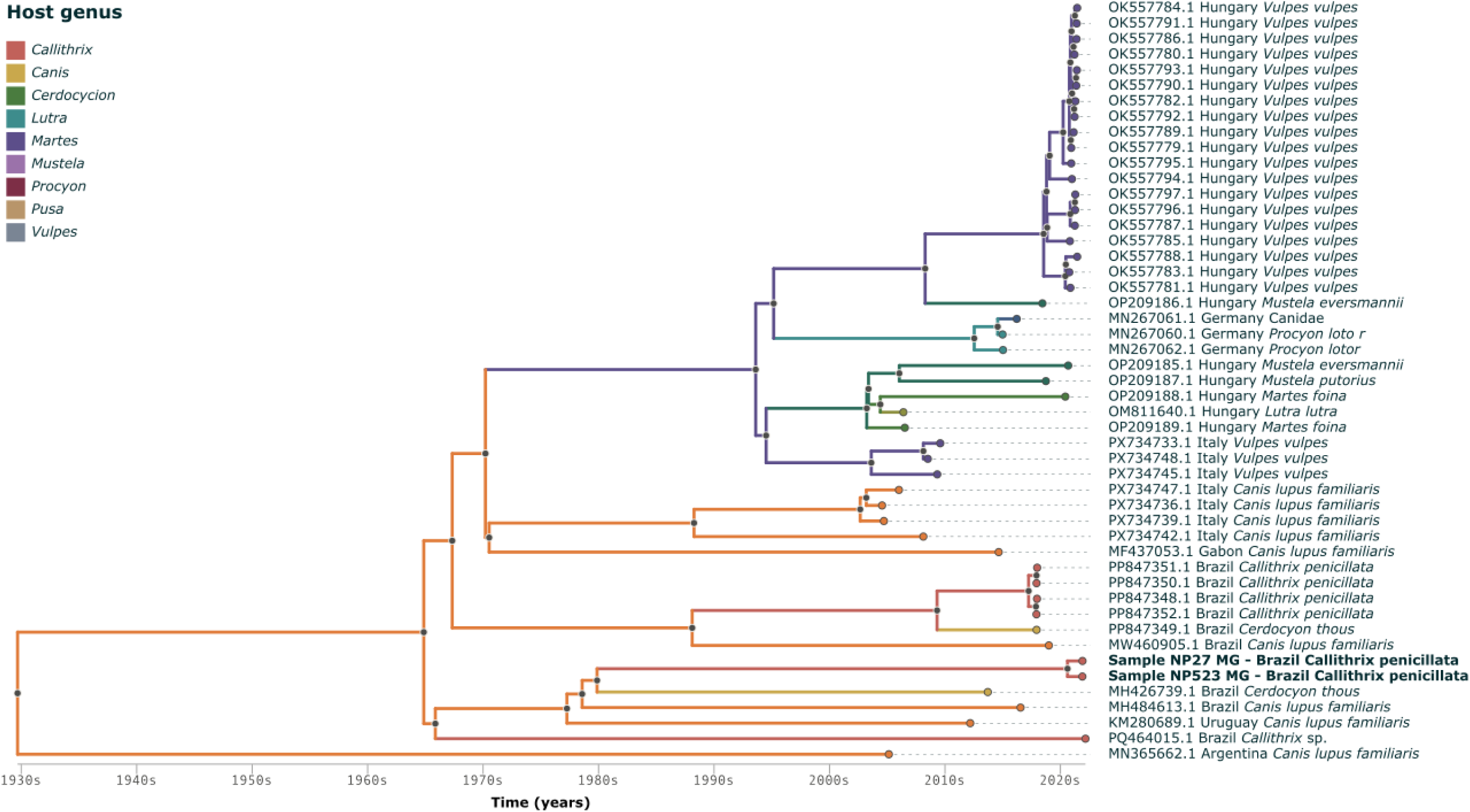
Bayesian time-scaled phylodynamic reconstruction of the European/South America 1 clade of Canine Distemper Virus (CDV). Host reconstruction using a nine-state model based on host genus (*Callithrix, Canis, Cerdocyon, Lutra, Martes, Mustela, Procyon, Pusa, and Vulpes*; DTA host model 2). Branches are colored according to the reconstructed ancestral host state at the genus level. Tip labels indicate GenBank accession, sampling location, host species, and collection date. The time scale (x-axis) is expressed in calendar years, with branch lengths proportional to time. The tree highlights host-switching patterns among the sampled host genera.

**Supplementary Figure S2.**
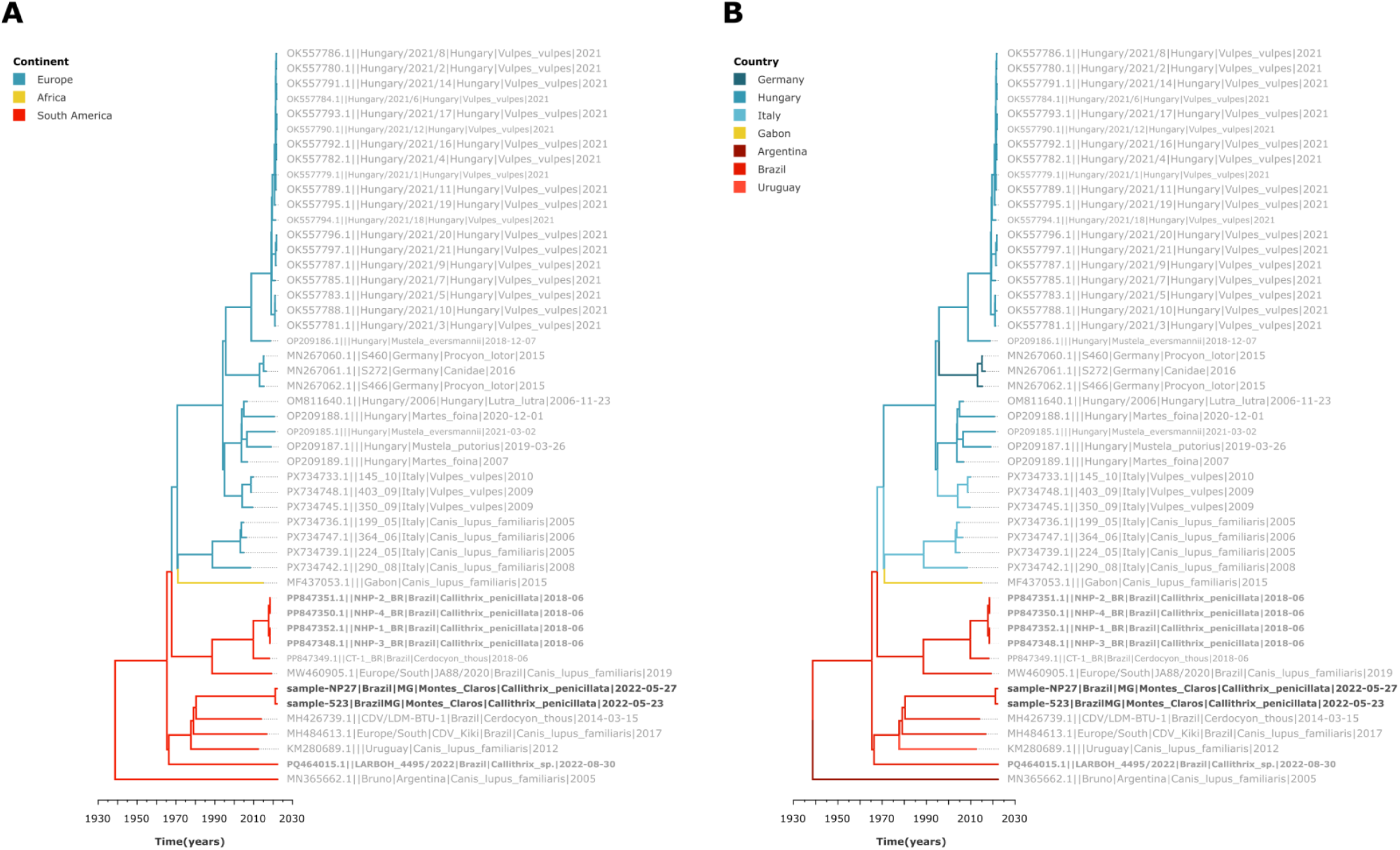
Bayesian time-scaled phylogeographic reconstruction of the European/South America 1 clade of Canine Distemper Virus (CDV). Maximum clade credibility (MCC) trees inferred from complete CDV genomes under a Bayesian molecular clock model, including the genomes generated in this study (bold). Branches are colored according to the reconstructed ancestral geographic state at the (A) continental and (B) country levels. Tip labels indicate GenBank accession, sampling location, host species, and collection date. The time scale (x-axis) is expressed in calendar years, with branch lengths proportional to time. The newly generated Brazilian genomes cluster within the South America 1 lineage, providing temporal and geographic context for their evolutionary relationships with previously reported CDV strains from Europe, Africa, and South America.

**Supplementary Figure S3.**
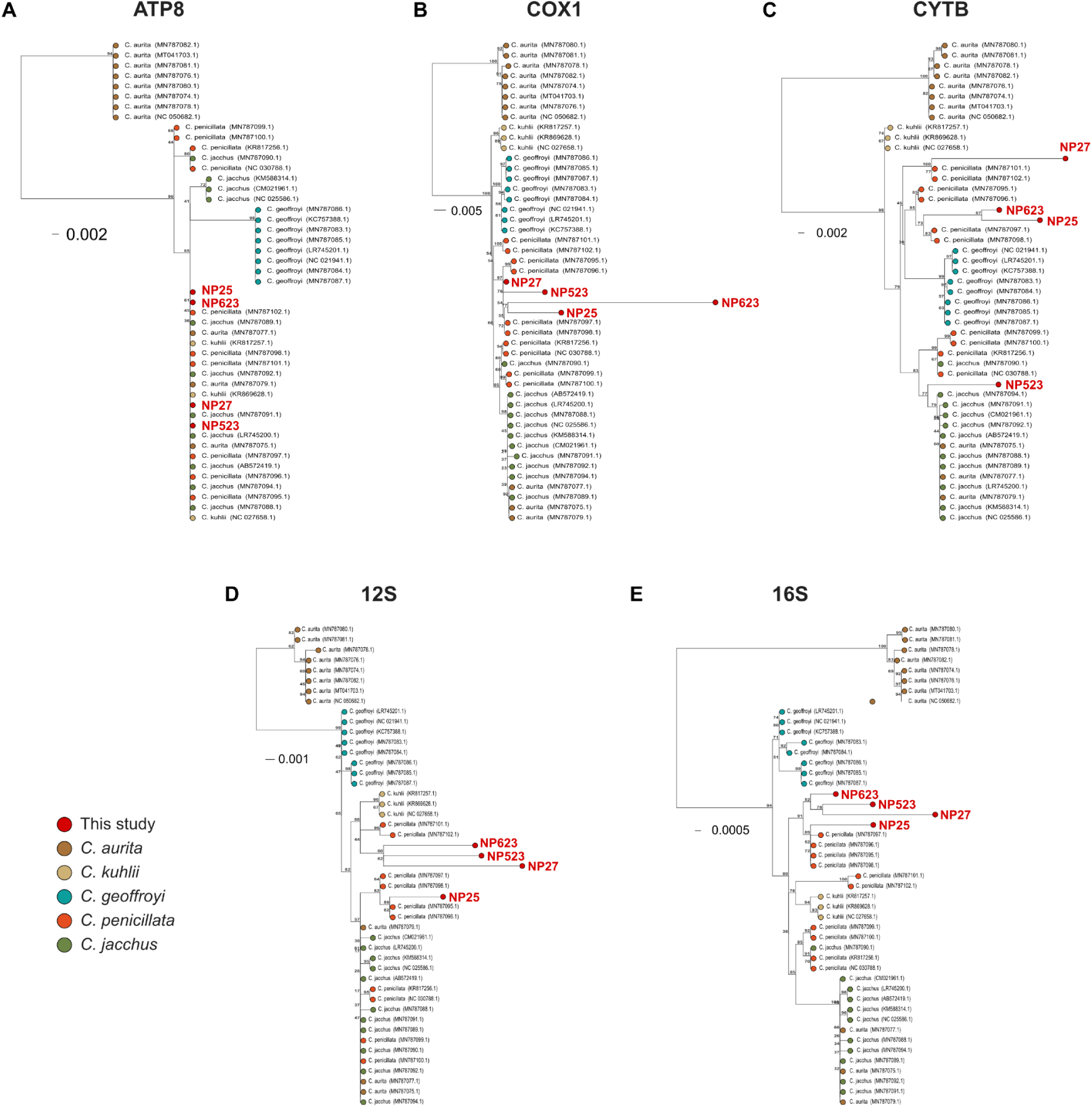
Gene-specific phylogenetic analyses of host mitochondrial sequences recovered from metagenomic data. Maximum likelihood phylogenetic trees inferred independently for the mitochondrial genes (A) ATP8, (B) COX1, (C) CYTB, (D) 12S rRNA, and (E) 16S rRNA. Sequences recovered in this study (red circles; NP25, NP27, NP523, and NP623) are shown together with representative Callithrix mitochondrial sequences from GenBank. Branch lengths are proportional to the number of nucleotide substitutions per site, with scale bars indicating evolutionary distance. While heterogeneous phylogenetic signal was detected across the trees inferred, the reconstructed phylogenies generally support the clustering of new sequences along *C. penicillata* clades, in agreement with the concatenated analysis.

**Supplementary Figure S4.**
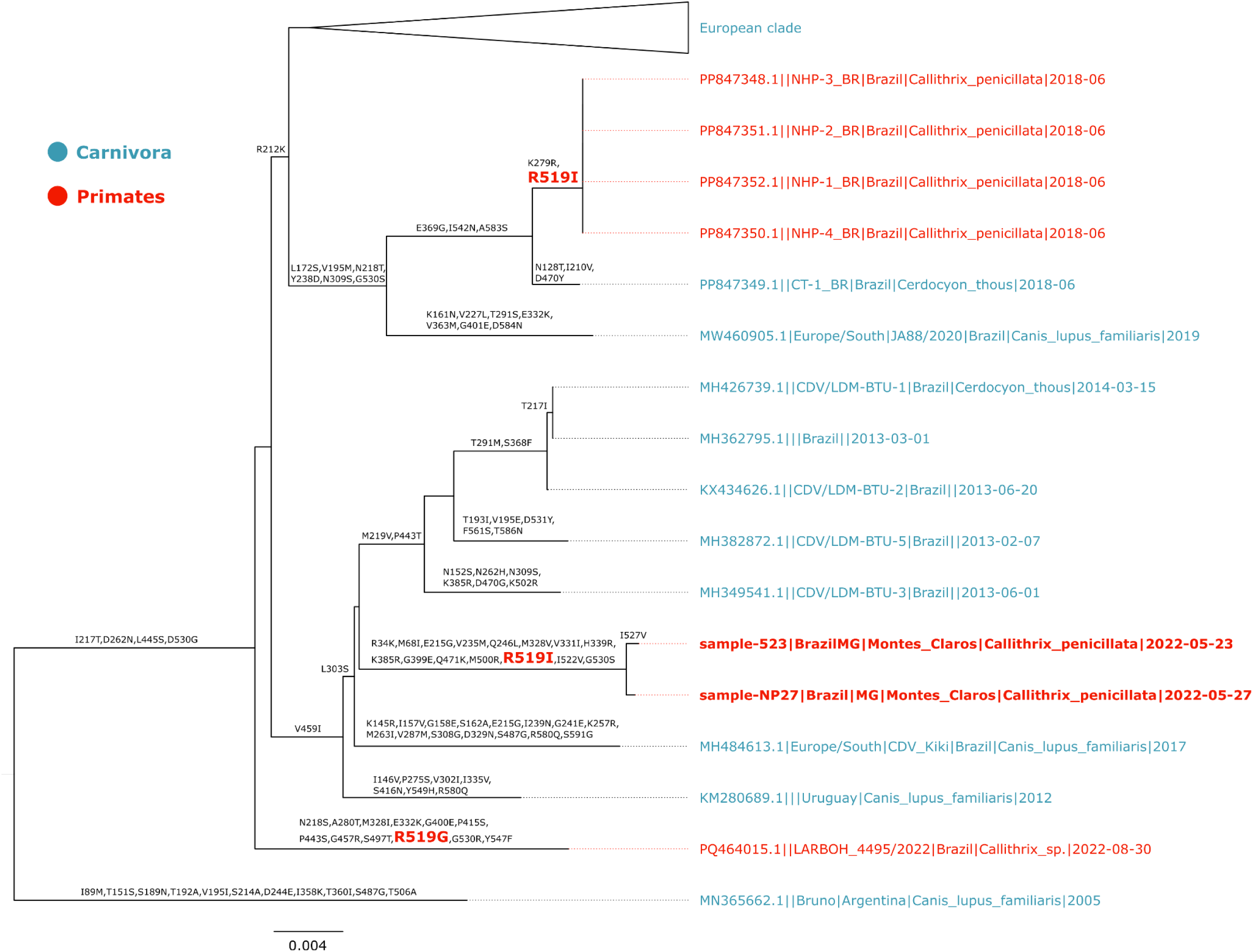
Maximum likelihood ancestral state reconstruction of amino acid substitutions at residue 519 of the CDV hemagglutinin (H) protein. Ancestral amino acid states were reconstructed on a maximum likelihood phylogeny of the European/South America 1 CDV clade to investigate the evolutionary history of residue 519 in the H protein. Tip labels indicate virus isolates, including accession number, sampling location, host species, and collection date. Branches and tip labels are colored according to host taxonomic order, with Carnivora shown in blue and Primates in red. Inferred amino acid substitutions at residue 519 are mapped onto the corresponding branches, highlighting independent occurrences of the R519I substitution in distinct primate-associated lineages and the R519G substitution in a separate primate-derived virus. These recurrent changes suggest convergent evolution at a residue might be implicated in host receptor interaction during CDV adaptation to non-carnivore hosts.

**Supplementary Figure S5.**
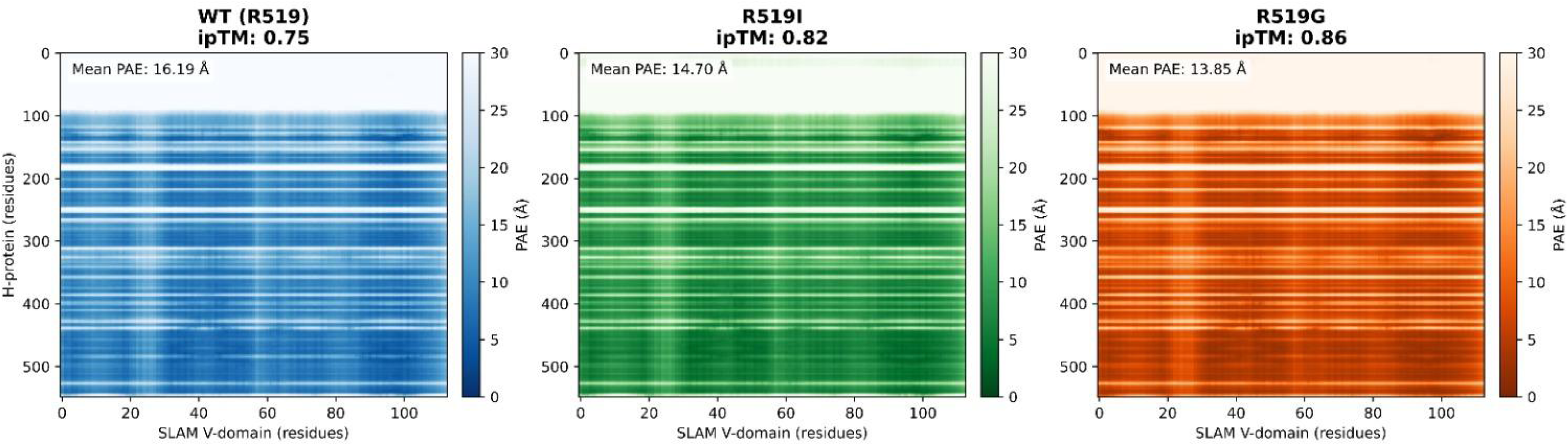
Predicted Aligned Error (PAE) heatmaps of the H-SLAM docking models. Interface PAE matrices (H-protein ectodomain, residues 59-607, against SLAM V-domain, residues 28-140) are shown for the top-ranked AlphaFold3 model of each variant: carnivore wild-type (WT, R519), primate R519I and primate R519G. The color scale represents PAE in Ångströms, from 0 to 30 Å. Darker shades indicate lower PAE and higher positional confidence, following the same color convention used by the AlphaFold Database. ipTM scores and mean interface PAE values are annotated in each panel. PAE values drop substantially across the receptor-binding head domain, including the region around residue 519.

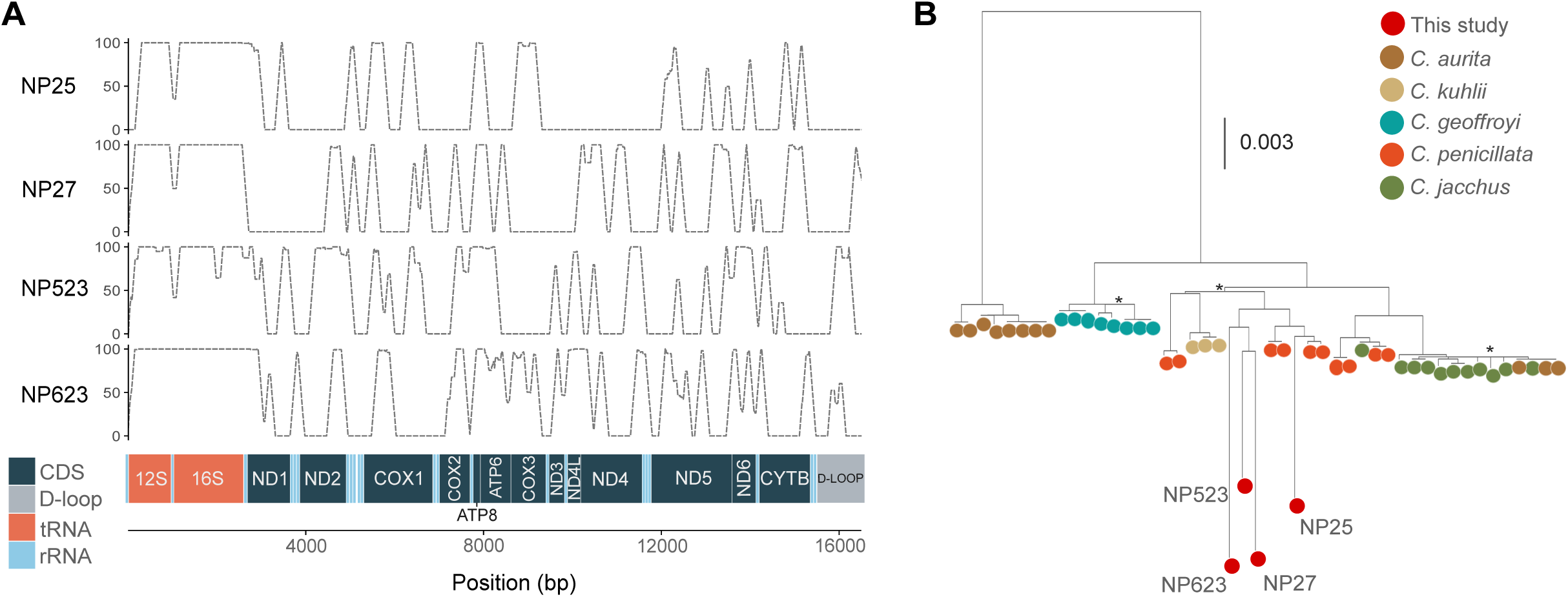

