## Supplementary figures and images for "Genomic Epidemiology of Canine Distemper Virus (CDV) in Neotropical Primates in Brazil Reveals Multiple Spillover Events and Convergent Evolution at Hemagglutinin Residue 519"

### Supplemental Figure S1

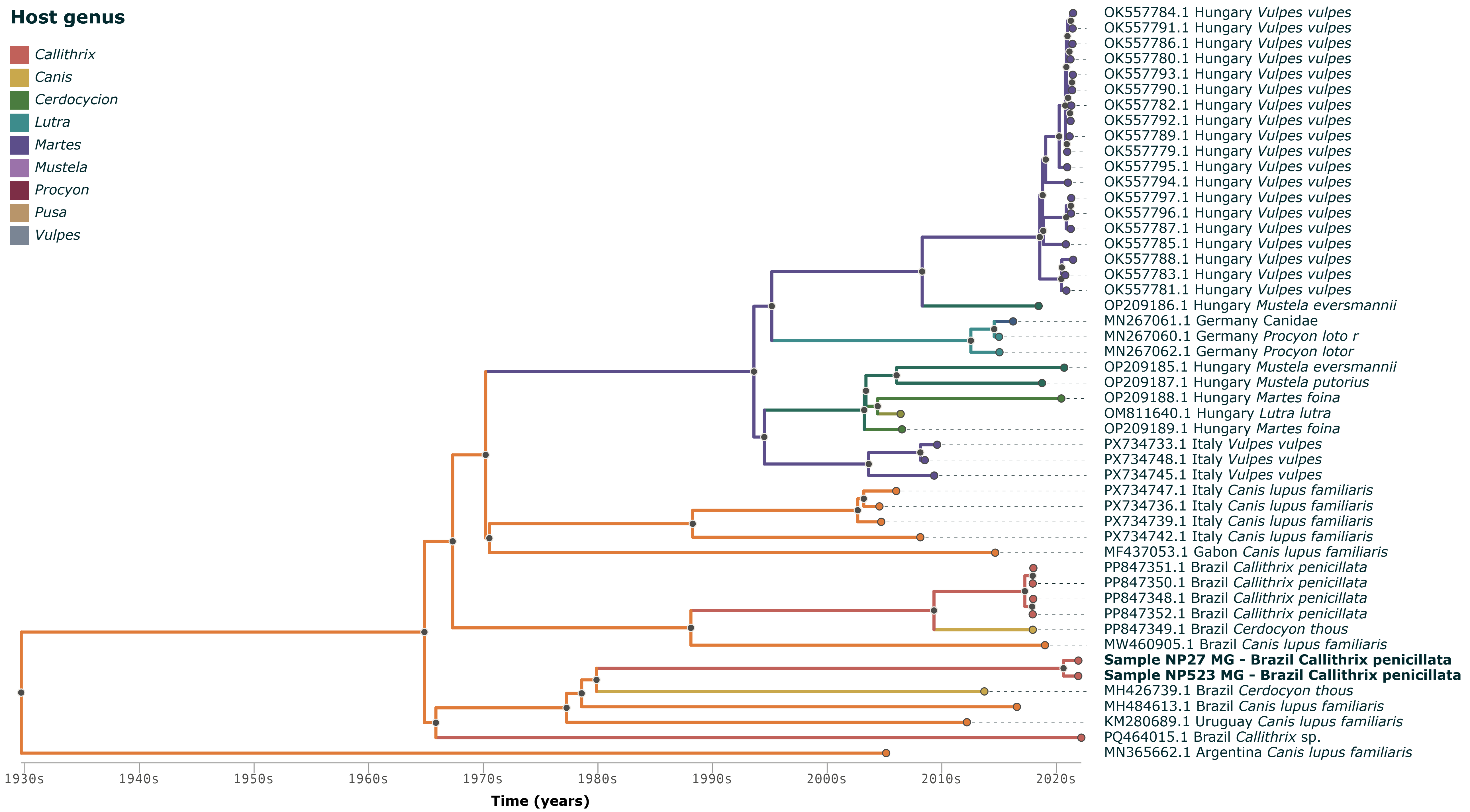

### Supplemental Figure S2

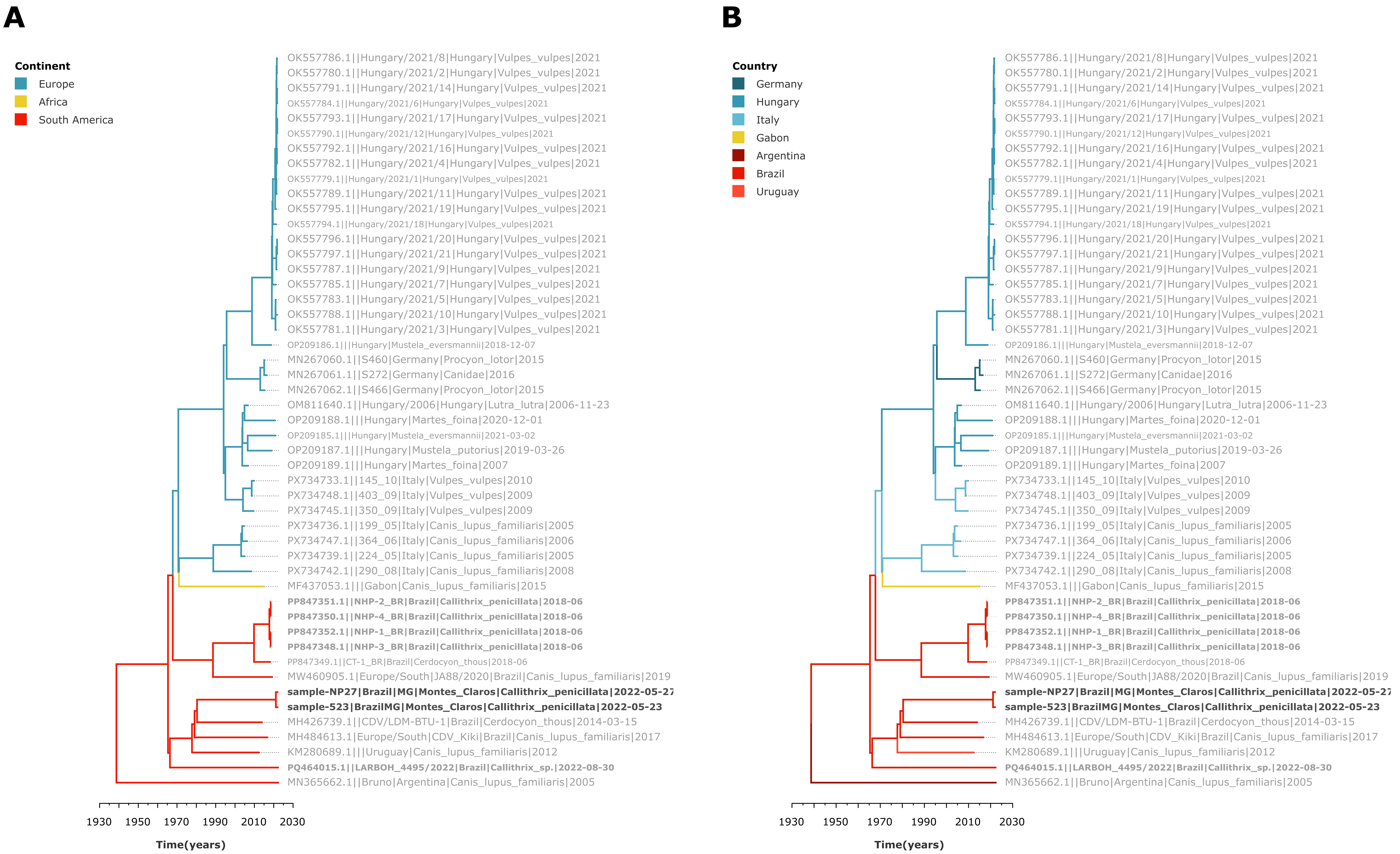

### Supplemental Figure S4

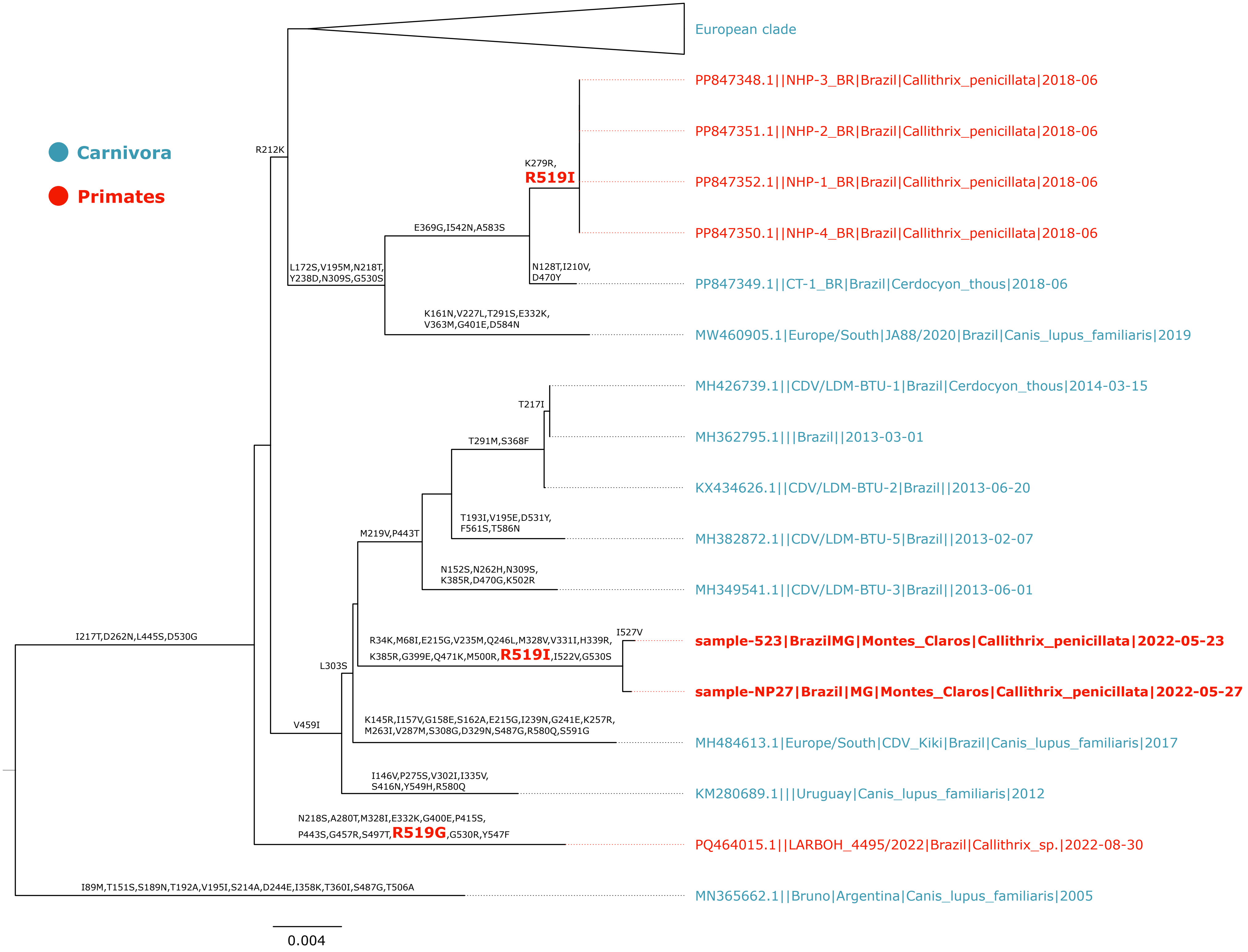

### Supplemental Figure S5

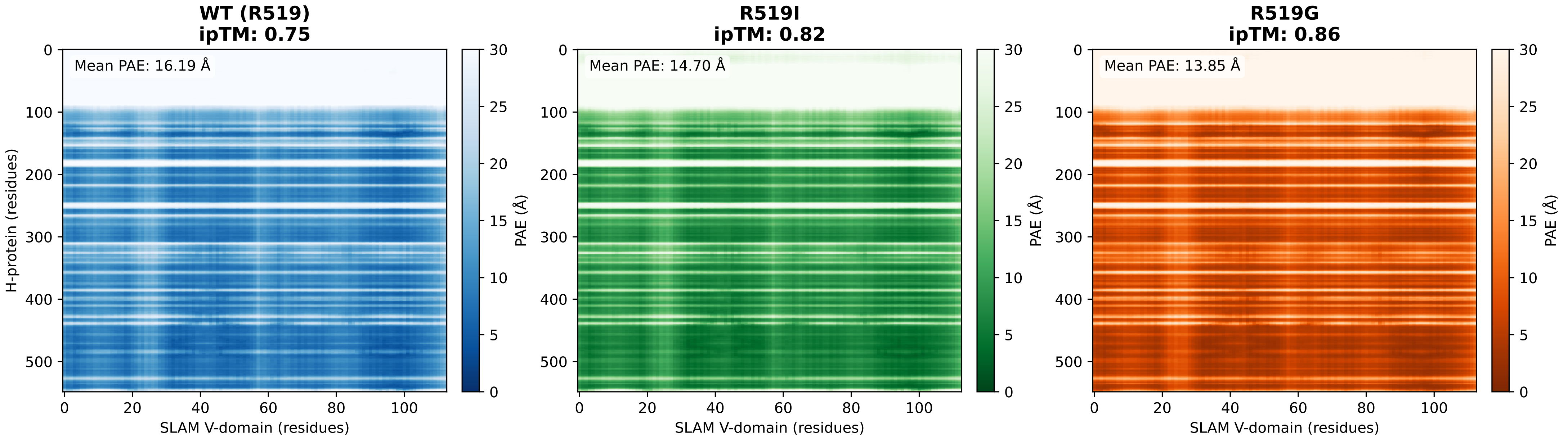
